# The effect of HNF4α knockout in beta cells is age and gender dependent

**DOI:** 10.1101/2024.12.16.628662

**Authors:** Catharina B.P. Villaca, Viviane R. Oliveira, Gustavo J. Santos, Fernanda Ortis

## Abstract

HNF4α is important for beta cells’ ability to adequately secrete insulin in response to glucose concentration and endoplasmic reticulum (ER) homeostasis. In humans, HNF4α mutation is responsible for Diabetes *mellitus* subtype MODY1, which has age determined onset. In addition, in other types of DM, there are evidences that gender can influence beta cell dysfunction, with possible involvement of ER stress pathways. Thus, we assessed the influence of gender and age on beta cell dysfunction induced by HNF4α absence. We used an animal model with specific beta cells KO for HNF4α, induced after birth (Ins. CRE HNF4α ^loxP/loxP^). Glucose intolerance is observed after 10 days of KO induction, at 50 days of age, with KO males (MKO) showing greater glucose intolerance than KO females (FKO). Percentage of insulin-positive cells in KO mice islets is lower compared to Control at all ages evaluated, with MKO having a lower percentage at later ages compared to FKO. Both KO groups have reduced beta cell mass and increased α-cell mass, which is higher in MKO. ER stress is induced in both KO groups. However, ER stress-mediated apoptosis is observed only in MKO. FKO shows evidence of beta cell differentiated state loss. Thus, loss of beta cells in HNF4α KO is influenced by gender and age, involves induction of ER stress, and is more pronounced in males, where ER stress-induced beta cell death is observed. Partial protection observed in females seems to involve dedifferentiation of beta cells.

## Introduction

Diabetes *mellitus* (DM) constitutes a spectrum of metabolic disorders marked by chronic hyperglycemia resulting from impaired insulin secretion or action (ADA, 2015). These disorders include polygenic forms, such as type 1 and type 2 DM, and monogenic forms (ADA 2015), such as the Mature Onset Diabetes of the Young (MODY), categorized into subtypes based on gene mutation leading to DM (Fajans 1989; Anık et al. 2015). MODY1 is a subtype defined by HNF4α mutation (Fajans 1989; Byrne et al. 1995, 1). Overall, HNF4α is an essential regulator of beta cell function (Stoffel and Duncan 1997; Ellard 2000, 4; Gupta et al. 2005; 2007; Barth et al. 2022).

The endoplasmic reticulum (ER) is crucial for the production of secreted proteins (Wei and Hendershot 1996), such as insulin (Dodson and Steiner 1998). During periods of high blood glucose, there is an increased demand for beta cells to produce and secrete insulin, resulting in an increased demand for ER folding capacity (Liu et al. 2018). This increase may lead to ER accumulation of misfolded proteins in its lumen, a condition known as ER stress (Arunagiri et al. 2018; Meyerovich et al. 2016). In this context, the Unfolded Protein Response (UPR) is activated, restoring ER homeostasis in most cases (Lenghel et al. 2020; Meyerovich et al. 2016). Thus, in beta cells, UPR activation allows adequate insulin secretion to match the demand, and its subsequent deactivation is important for cellular stress recovery (Xin et al. 2018).

Although the activation of the UPR in a cyclic manner is beneficial for beta cells’ secretory capacity, its chronic activation, as observed during chronic hyperglycemia or insulitis, can switch to a pro-apoptotic profile (Decio L. Eizirik and Cnop 2010; Oslowski and Urano 2010; Meyrovich et al, 2016). The chronification of the ER stress leads to beta cell apoptosis, further impairing beta cell population ability to reestablish normoglycemia through insulin secretion (Le May et al. 2006; Chan et al. 2015; Hu et al. 2019). ER stress-mediated apoptosis is an important component of DM development (Décio L. Eizirik, Cardozo, and Cnop 2008; Meyrovich et al, 2016). There are evidences that HNF4α may modulate UPR pathways in beta cells (Sato et al. 2012; Moore et al. 2016). However, these findings disagree on whether HNF4α contributes to a pro-adaptative or a pro-apoptotic profile of UPR.

This conflict can be in part due to the model used since there are other influences on beta cell function, such as the environment of the pancreatic islets (Brereton et al. 2015a) as well as gender and age (Le May et al. 2006; Aguayo-Mazzucato et al. 2017). For instance, estrogen has a protective effect on the beta cell apoptosis mediated by ER stress, and older females lose this protection due to menopause, having similar outcomes as males (Basu, Dube, and Basu 2017).

In addition to apoptosis, it has been established that beta cell failure in DM may also occur by loss of differentiated status (Patel and Remedi 2024; Bensellam, Jonas, and Laybutt 2018), as observed for glucotoxicity that affects oxidative stress, inflammation, and ER stress pathways (Khin, Lee, and Jun 2021) or direct induction of ER stress (Zhang et al. 2024) leads to the loss of beta cell mature phenotype.

Here, in a mouse model of HNF4α specific beta cell knockout (KO), we investigate the influence of gender and age on beta cell UPR activation and dedifferentiation in the absence of HNF4α.

## Materials and methods

### Ethical approval

Animal experiments conducted in this study were in accordance with local law and approved by the Animal Experimentation Ethics Committee (CEUA) of the Biomedical Sciences Institute (ICB) of the University of Sao Paulo, Brazil, under protocol number 9180111119.

### Animals

#### Origin, welfare, and breeding

InsCre HNF4α^loxP/loxP^ mice from "Biotério Setorial do Departamento de Biologia Estrutural e Funcional da Universidade Estadual de Campinas/SP" were adapted to and maintained in the animal housing facilities of the Department of Cell and Developmental Biology (ICB-USP). Mice had access to food (Nuvilab cr1, Nuvital Nutrientes LDTA, Curitiba) and water *ad libitum* and were maintained at a controlled temperature (22 °C) in a 12-12 h light/dark cycle and maintained in collective cages (max 4/cage).

InsCre ^+/+^ and InsCre ^-/-^ were the experimental groups, respectively knockout (KO) and control (Ctr). Mice were genotyped using primer sequences recommended by The Jackson Laboratories (Maine, USA).

#### Ages of interest

For the evaluation of islet cells, we selected 50 days of age (10 days after HNF4α KO induction), 90, and 150 days of age (Figure 1), equivalent to early adulthood in humans. In this period, beta cell dysfunction onset due to MODY1 is associated with hyperglycemia (Fajans 1989).

**Figure 1:**
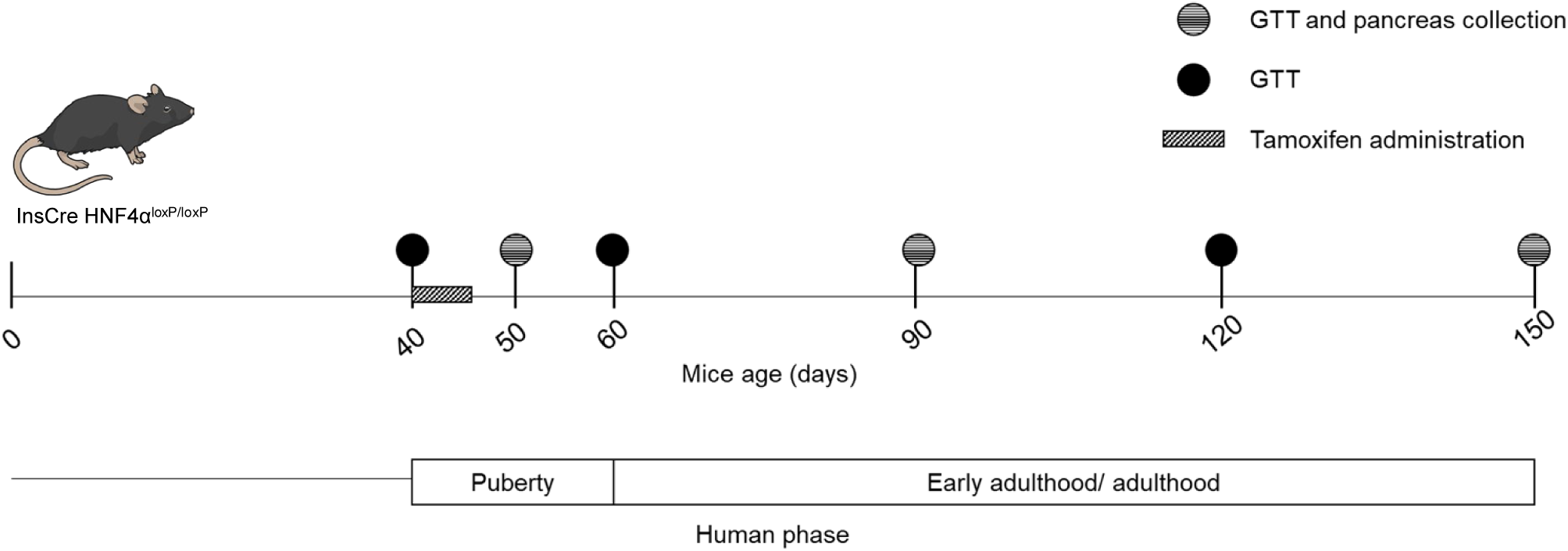
Experimental design. Scheme highlighting key mice ages selected for the study and the experiments performed and its correlation with human ages. GTT: Glucose Tolerance test.

As age itself might affect glucose handling (Hoffman et al. 2000), we also evaluated glucose tolerance at 40 days (before induction of the KO), 60 days (final stages of puberty), and 120 days (midpoint) of age in addition to 50, 120 and 150 days of age (Figure 1). We weighed mice weekly on the same day and time of the glucose tolerance test (GTT) assay.

#### Tamoxifen administration for Knockout induction

Mice from both experimental groups (KO and Ctr) were treated with tamoxifen (#T5648, Sigma-Aldrich) diluted in sesame oil (#3547-250, Sigma-Aldrich), for 5 days (75 mg/Kg) (Figure 1) as previously described (Moore et al. 2016).

#### Glucose tolerance test (GTT)

Animals were fasted overnight for 10-12 hours, and fasting blood glucose (time 0) was evaluated through a blood sample obtained from the caudal vein measured in an Accu-Check glucometer (Roche). Then, glucose solution (1.25 g glucose/kg of animal weight) was administered intraperitoneally, and blood glucose was evaluated at 15, 30, 60, 90, and 120 min. This test was performed on animals at 40, 50, 60, 90, 120, and 150 days of age (Figure 1).

### Histological analysis

#### Pancreas processing and hematoxylin and eosin staining

At 50, 90, and 150 days of age, the pancreata were collected and processed for histological assays, sectioned, and stained by hematoxylin and eosin conducted as previously described (Villaça et al. 2021).

#### Immunohistochemistry

The tissues were prepped for immunohistochemistry assays as previously described (Villaça et al. 2021), and primary and secondary antibodies were diluted in TBS+BSA 3%. The stain was developed with Diamino-Benzidine (DAB) and hydrogen peroxide (Synth) against the stain with Mayer’s Hematoxylin and assembled in Damar gum. Primary antibodies used were anti-HNF4α (1:50, C11F12, Cell Signaling Technology), anti-insulin (1:100, sc-9168, Santa Cruz) and anti-glucagon (1: 200, L16G, Immunity Biotechnology), secondary antibody used was horseradish peroxidase-conjugated anti-rabbit IgG (1:200; MJ163713; Thermo Fisher Scientific; Waltham, MA, USA) or anti-mouse IgG (1:75; 31430; Santa Cruz Biotechnology).

#### Immunofluorescence

Tissue sections were processed as previously described (Villaça et al. 2021). The mounting was done with Fluoroshield with DAPI (F6057; Sigma Aldrich). Slides were stored in the freezer for a maximum of 1 week before microscopic photography.

Slides were immunostained with anti-CHOP (1:100, Cell Signaling Technology), anti-XBP1-s (:50, 40435S, Cell Signaling Technology), anti-GRP78 (1:20, PA1104A, Invitrogen), anti-Glut-2 (1:10, BS-10379R, ThermoFisher), anti-PAX-4 (1:50, PA1108, Invitrogen), anti-NGN3 (1:100, BS −0922R, Invitrogen) or anti-SOX9 (1:150, PA581966, Invitrogen) in combination with anti-insulin (1:100, sc-9168, Santa Cruz or 1:100, 53-9769-82, ThermoFisher) and corresponding secondary antibody anti-mouse (1:100, A1100 Invitrogen) or anti-rabbit (1:100, A11036 Invitrogen).

#### Islet image acquisition and quantification

All slides were documented using photomicrographs with Zeiss Axioskop 2. HE staining was documented with a 5X magnification objective to determine the section area and 40X to determine the islet area, circularity, and the relative number of nuclei per islet. The immunostains were photographed at 40X magnification.

Slides and the images were independently evaluated by two individuals, one aware and one unaware of sample identity. The agreement between the findings obtained by the two observers was at least 90%. Quantification was performed using the "QuPath" software (Bankhead et al. 2017)).

The islets were manually traced, creating Regions Of Interest (ROI). Images of islets not used for this study were used to determine a positive threshold value based on stain intensity for each of the stains of interest to determine positive labeling. For the HNF4α, the threshold value was 0.75, insulin was 0.35, and glucagon 0.55. Within the islets ROI we utilized the predetermined intensity value to determine individual islet positivity to the target of interest. Additionally, for the insulin labeling, we measured the variation of insulin intensity labeling, as this measurement serves as a proxy for protein abundance, as previously described (Braun et al., 2013; Meyerholz and Beck, 2018). This was calculated using the formula: H-Score=3*intense score+2*medium score+1*weak score, as previously established (Bankhead et al. 2018).

From the quantification of alpha and beta cell masses, we calculated using the formula: mass= (total positive area of the marker)/ (total area of the cut) x wet weight of the pancreas (modified from Dos Santos et al. 2021).

The QuPath software ("object classification") quantified the co-localization with insulin immunofluorescence assays. This function allows the evaluation of cell by the cell of positivity for insulin labeling and if this is in combination with positivity for other proteins (CHOP, GRP78/BiP, XBP1-s, GLUT-2, PAX-4, NGN3, and SOX9). The data was expressed as the percentage of positive cells for the label of interest in relation to the total of positive cells for insulin.

#### Statistical analysis

Statistical analysis was performed using the GraphPad Prism v9 software. Results were analyzed by two-way ANOVA with multiple comparisons (Tukey post-test). Except for the area under the curve (AUC) of the GTTs analysis, a three-way ANOVA with multiple comparisons was used.

## Results

### Animal model characterization

The percentage of islet cells positive for HNF4α immunostaining was reduced, around 80%, in KO mice compared to Ctr, five days after the last tamoxifen administration, as previously described (Gupta et al. 2005). It is important to note that there was no difference between genders, with equal efficiency of KO in both females and males (Supplementary Figure 1). The HNF4α KO body weight was not affected during the analyzed period. As expected, males had significantly higher body weight gain (Reed, Bachmanov, and Tordoff 2007) (Figure 2A). Females, independent of the genotype, had significantly higher proportional pancreas weights at all ages when compared to males (Figure 2B). Comparison between genotypes showed that KO had a higher proportional pancreas weight after 90 days compared to Ctr (Figure 2B).

**Figure 2:**
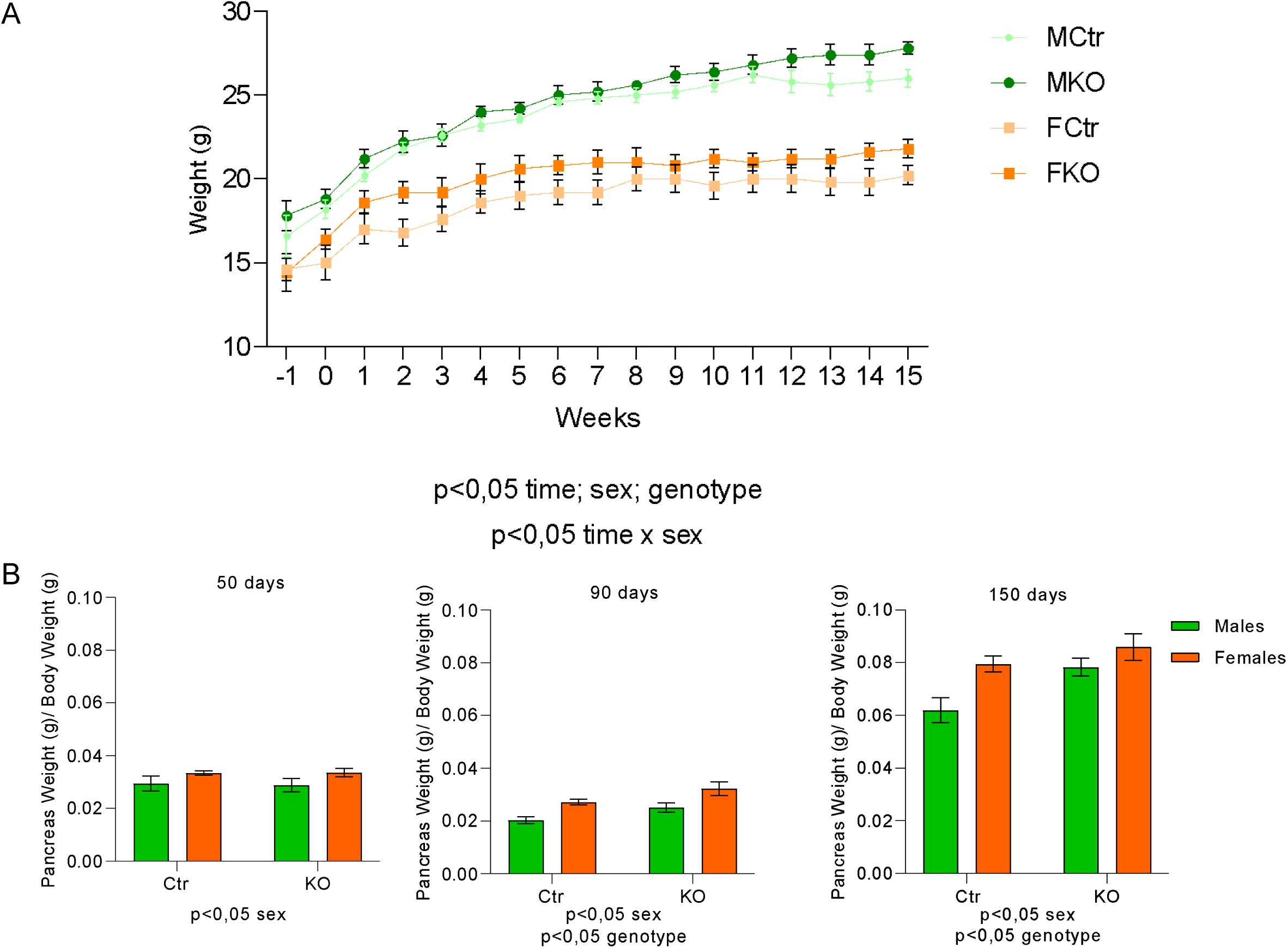
Body and pancreas weight. Weight gain during experimentation phase (week 0 being the tamoxifen administration week) (A) and pancreas wet weight after dissection (B) at 50, 90 and 150 days of age. Source of variation is reported under the graph (n=5/ group), data is shown as mean± SEM

At 40 days of age, before KO induction, all groups had similar GTT curves (Figure 3A). Ten days after KO induction (50 days of age), KO animals of both genders presented increased glycaemia, 15 minutes after glucose administration, compared to the respective Ctr. At 30 minutes, only KO males (MKO) maintained this difference (Figure 3A). At a later age in puberty (60 days), only MKO presents higher glycemia at the first 3 points of the curve. Although, at this age, Ctr females (FCtr) present higher blood glucose at 15 min, when compared to males of the same genotype, there is no difference between the KO groups (Figure 3A).

**Figure 3:**
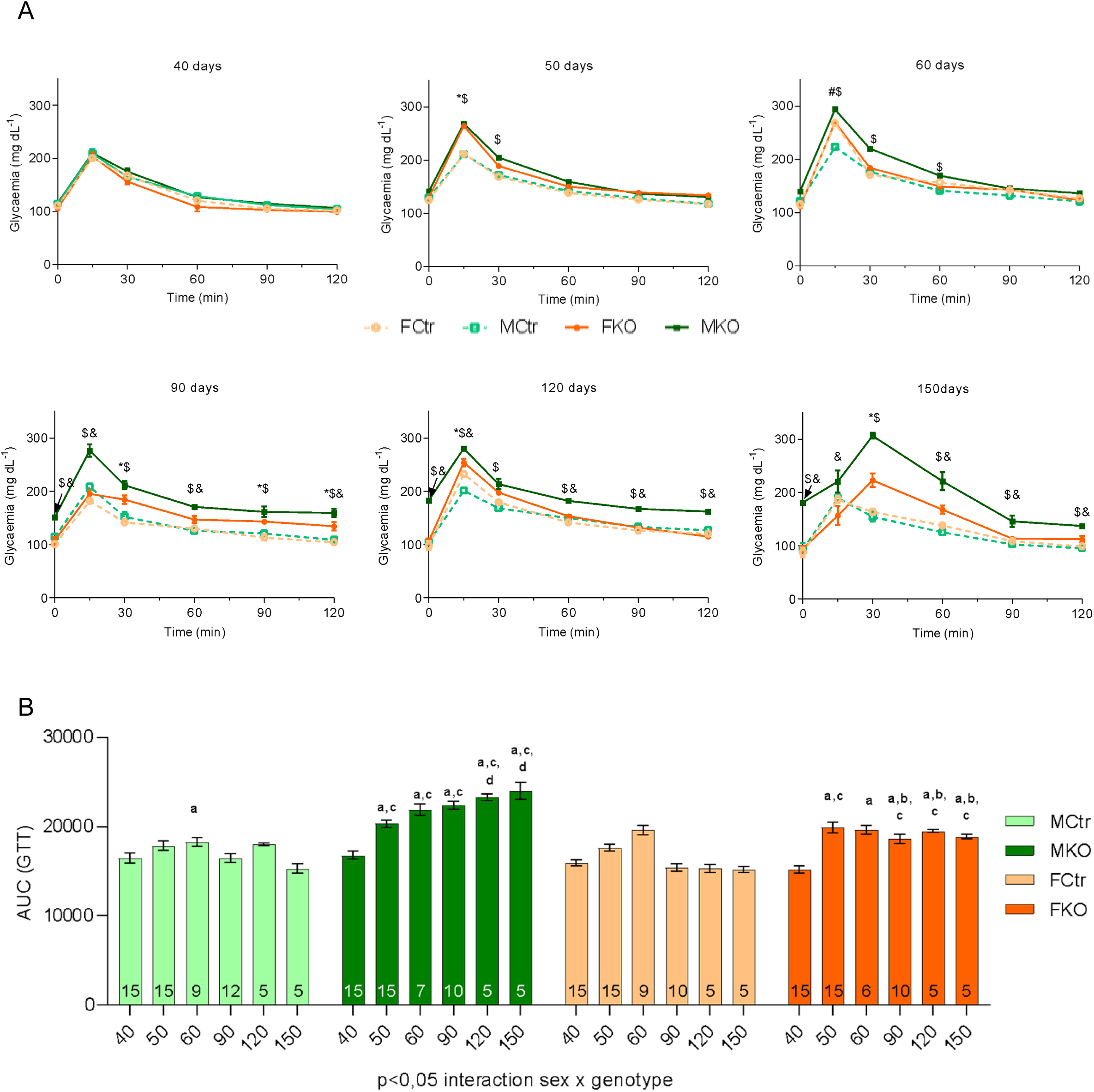
Glucose tolerance test. Glucose tolerance test at 40, 50, 60, 90, 120 and 150 days of age from all groups: FCtr (Female Control); MCtr (Male Control); FKO (Female KO) and MKO (Male KO). p<0,05: * FKO vs FCtr, $ MKO vs MCtr, & MKO vs FKO, #MCtr vs FCtr (A). Area under the curve of these tests P<0,05: a vs 40 days old (same group), b p<0,05 vs male (same age and genotype), c vs Ctr (same age and gender), d p<0,05 vs 50 days old (same group) (B). Source of variation is reported under the graph. N of each test is reported within the bars in B, data is shown as mean± SEM

At the beginning of the young adult phase (90 days of age), MKO showed hyperglycemia at all points of the GTT, as well in fasting, compared to the MCtr. In contrast, the KO females (FKO) only showed this increase in later points (after 90 min) (Figure 3A). This glucose intolerance observed in all time points evaluated for MKO persists at 120 days of age, still in the young adult phase, in contrast, FKO at 120 days of age present higher glycemia only at 15 min (Figure 3A). In the final part of the young adult phase (at 150 days of age), there is a change in glucose handling for KO animals since the GTT curve peak that occurs typically at 15 min is observed at 30 min in this group, independent of the gender. The curve remains the same in Ctr animals, demonstrating that the temporal change occurs in response to the absence of HNF4α (Figure 3A).

Next, we evaluate the temporal progression of gender impact in animals of different genotypes using the 3-factor analysis of the AUC to assess the effect of gender on glucose tolerance in function of knockout and time. This showed that MKO has reduced glucose tolerance compared to FKO after 60 days of age (Figure 3B). In addition, the MKO shows a progressive loss in glucose tolerance in function of the age, which is not seen in the FKO (Figure 3B)

### Islet structure and cell populations

Qualitative analysis of islet histology from KO and control animals showed decrease in islet area of KO animals at later ages (Figure 4A). Quantification of these parameters showed that a decrease in area (Figure 4B) and circularity (Figure 4C) in MKO animals is absent at 50 days and only observed at later ages, 90 and 150-day-old animals, compared to both MCtr and FKO. For FKO, however, only at 150 days of age there was a reduction of islet area compared to FCtr (Figure 4B), with no changes in islet circularity (Figure 4C). In agreement with these results, reduction of islet cell number, evaluated by number of nuclei/islet (Figure 4D), is observed in MKO islet when compared to MCtr at 90 and 150 days of age and only at 150 days when compared to FKO (Figure 4D). Thus, the influence of gender (P<0.05 gender x interaction genotype) in the effect of HNF4α KO in islet area and circularity is observed already at 90 days of age (Figure 4).

**Figure 4:**
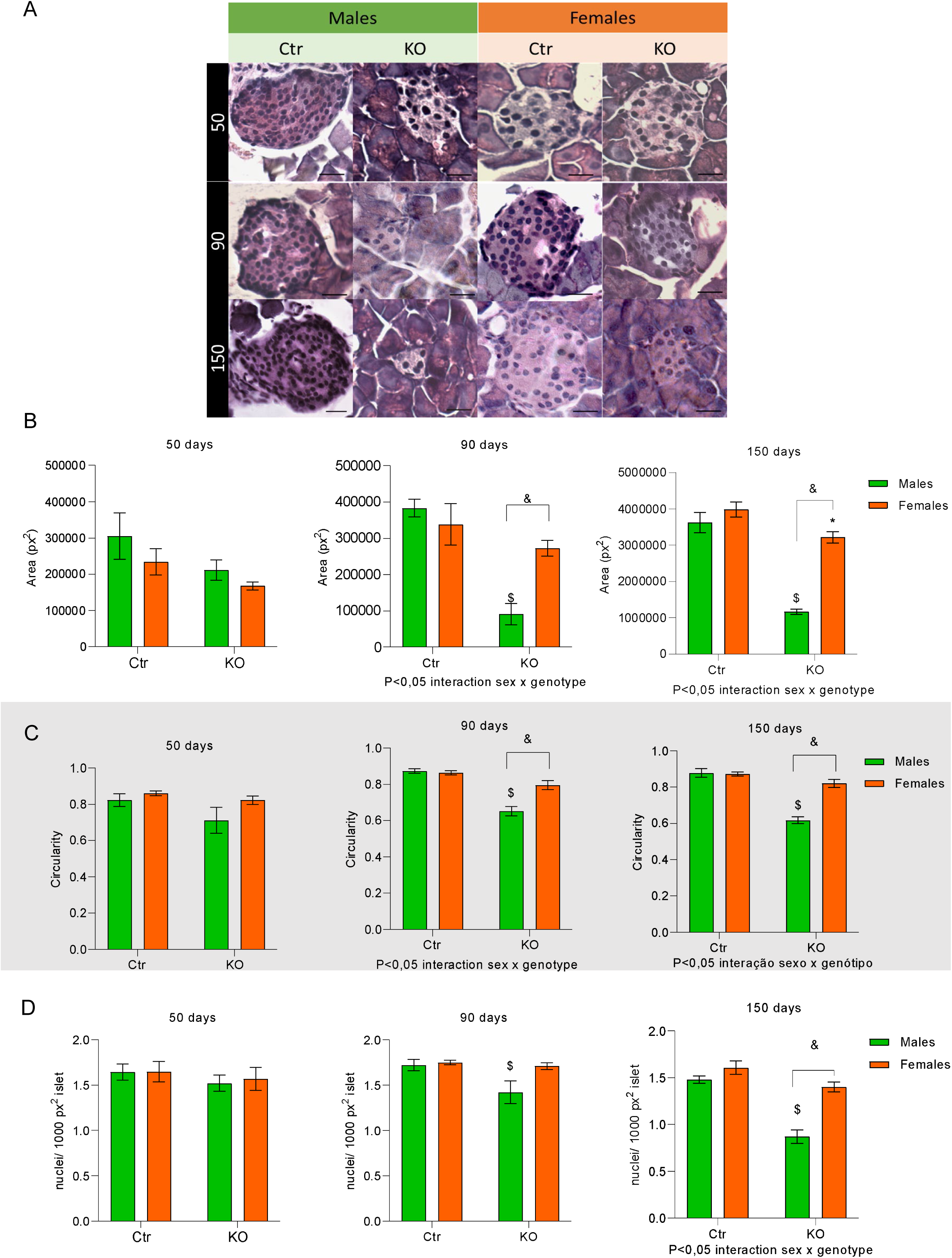
Islet morphometric parameters. Representative images of all 12 groups:4 genotypes FCtr (Female Control); MCtr (Male Control); FKO (Female KO) and MKO (Male KO) at 3 different ages (A). Quantification of average islet’s area (B), circularity (C) and nuclei per islet (D). N=5/group, data is shown as mean± SEM, p<0,05: *FKO vs FCtr, $ MKO vs MCtr and & FKO vs MKO, source of variation is reported under the graph. Scale bar: 50 μm.

In both female and male control mice at all ages evaluated, insulin immunostaining is located in the center of the islet (Figure 5A) and glucagon in the periphery (Figure 5B), according to the expected cell distribution in rodents (Kim et al. 2009). In KO animals at 50, 90 and 150 days of age, insulin immunostaining is less evident, as shown by a decrease in the percentage of insulin-positive islet area, already at 50 days of age, compared to Ctr (Figure 6A). This loss of insulin-positive area in islets from KO animals is progressive with age, more evident at 90 and 150 days. This progressive reduction is more intense in MKO, going from 37% insulin positive islet cells at 50 days to 9% at 150 days of age, compared to FKO, going from 39% insulin positive islet cells at 50 days to 24% at 150 days of age (Figure 5 and 6A). The intensity of insulin labeling (H-Score) is also significantly reduced in KO animals compared to Ctr at all ages, regardless the gender (Figure 6B). Finally, beta cell mass is reduced in KO animals compared to the respective Ctr from 90 days of age, regardless the gender (Figure 6C).

**Figure 5:**
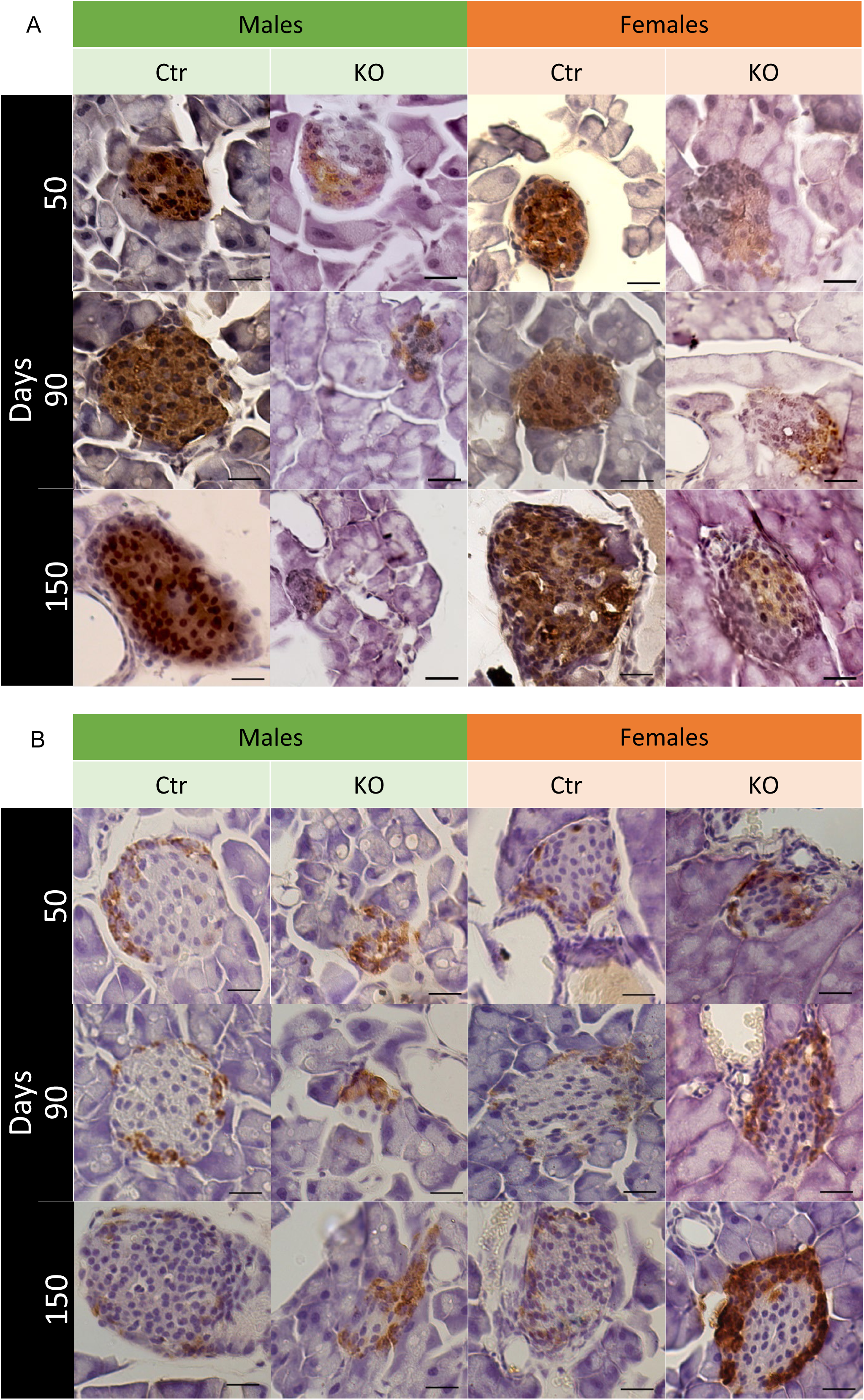
Insulin and glucagon stains. Representative images of all 12 groups:4 genotypes FCtr (Female Control); MCtr (Male Control); FKO (Female KO) and MKO (Male KO) at 3 different ages, showing the localization of the immunostaining and intensity of insulin (A) and glucagon (B). Scale bar: 50 μm.

**Figure 6:**
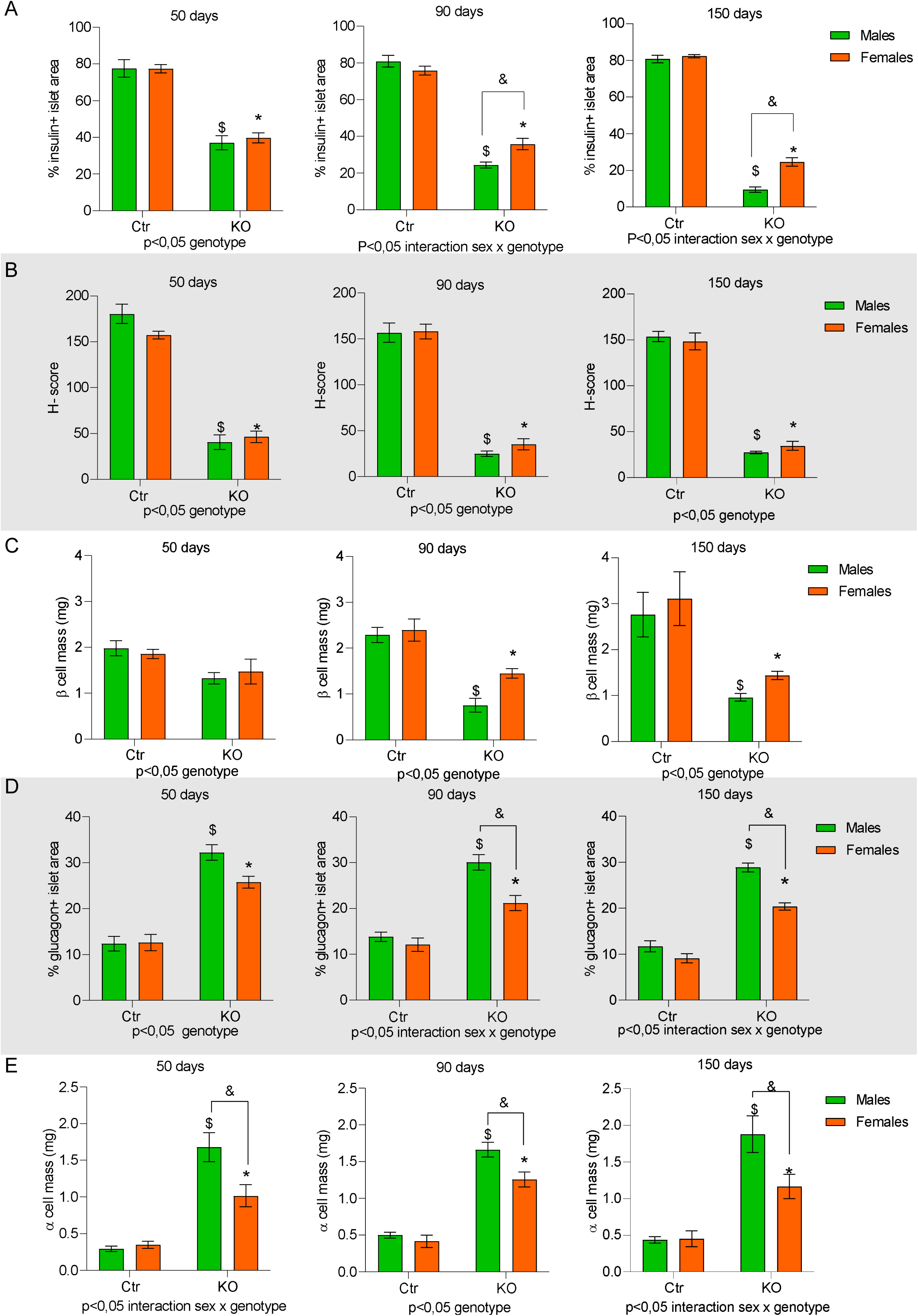
Quantification of the insulin and glucagon immunostain. Quantification of all 12 groups:4 genotypes FCtr (Female Control); MCtr (Male Control); FKO (Female KO) and MKO (Male KO) at 3 different ages for insulin area (A), h-score (B), beta cell mass (C), glucagon area (D) and alpha cell mass (E). N=5 group, data is shown as mean± SEM, p<0,05: *FKO vs FCtr, $ MKO vs MCtr and & FKO vs MKO, source of variation is reported under the graph.

Glucagon immunostaining is more prominent in the KO than in the Ctr group (Figure 5B). The percentage of glucagon-positive cells per islet area is increased in KO groups compared to control animals already at 50 days of age. From 90 days of age, this parameter starts to be influenced by gender, with MKO showing a significant increase in glucagon-positive cells compared to females of the same genotype (Figure 6D). Finally, the alpha cell mass is higher in the KO groups compared to the respective Ctrs at all ages, with MKO having a significant increase compared to FKO (Figure 6E).

### Endoplasmic reticulum stress markers in insulin-positive cells

The co-localization of CHOP (green) with insulin (red) was exacerbated in islet cells from MKO animals at all ages analyzed, as highlighted in the right panel (Figure 7A). The quantification of this co-localization confirms that only in MKO animals the absence of HNF4α leads to an increase in CHOP in beta cells (Figure 8A).

**Figure 7:**
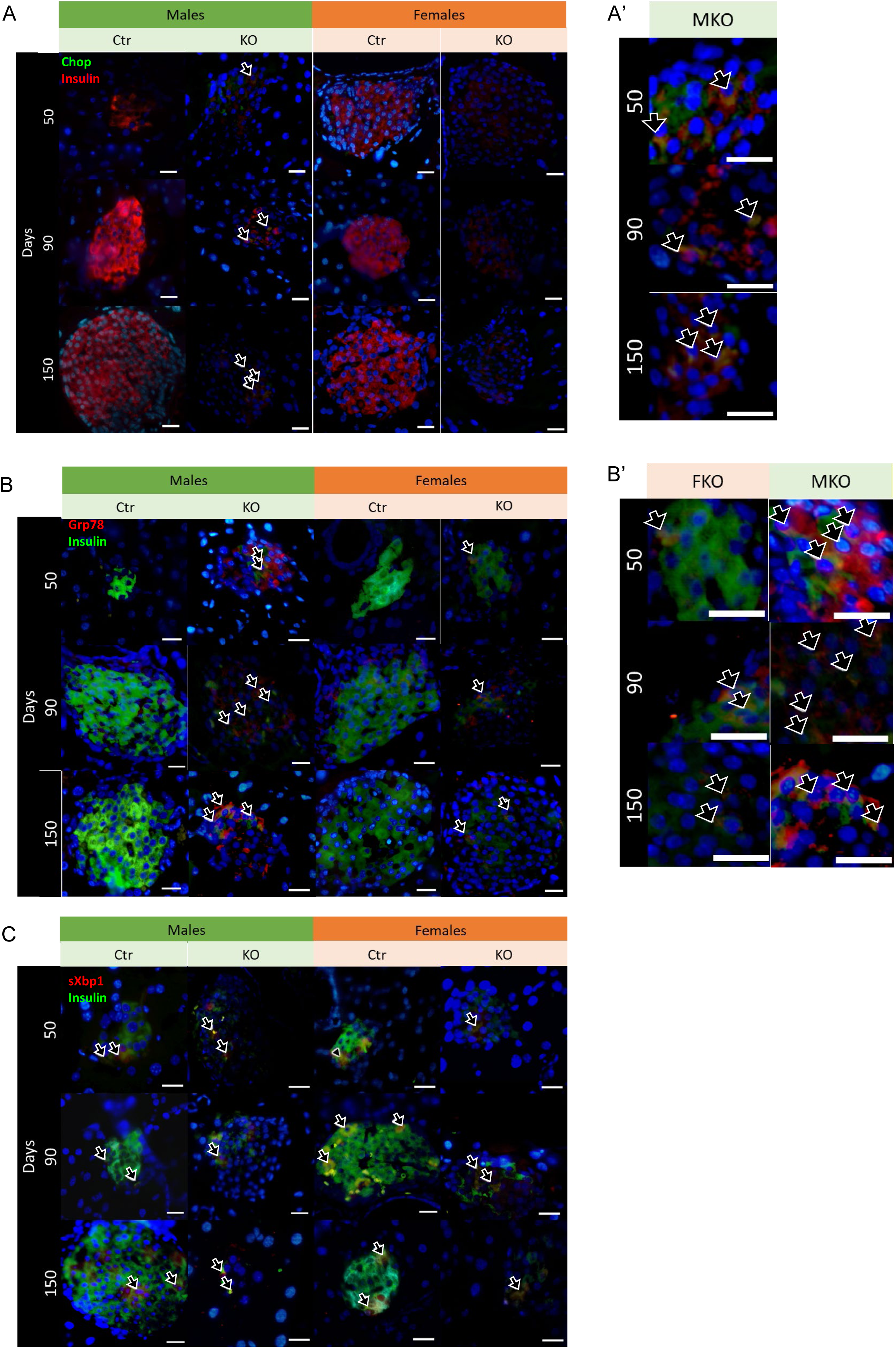
Endoplasmic reticulum stress markers colocalized with insulin. Representative images of all 12 groups:4 genotypes FCtr (Female Control); MCtr (Male Control); FKO (Female KO) and MKO (Male KO) at 3 different ages, immunofluorescence showing the co-localization (yellow) of insulin (red) with CHOP (green) (A); or insulin (green) with GRP78 (red) (B) or XBP1-s (red) (C). On the right side of the panels there is a zoomed representative image of the co-localization observed in A (A’) and B (’B). Nuclei were stained with DAPI (blue). Scale bar: 50 μm.

**Figure 8:**
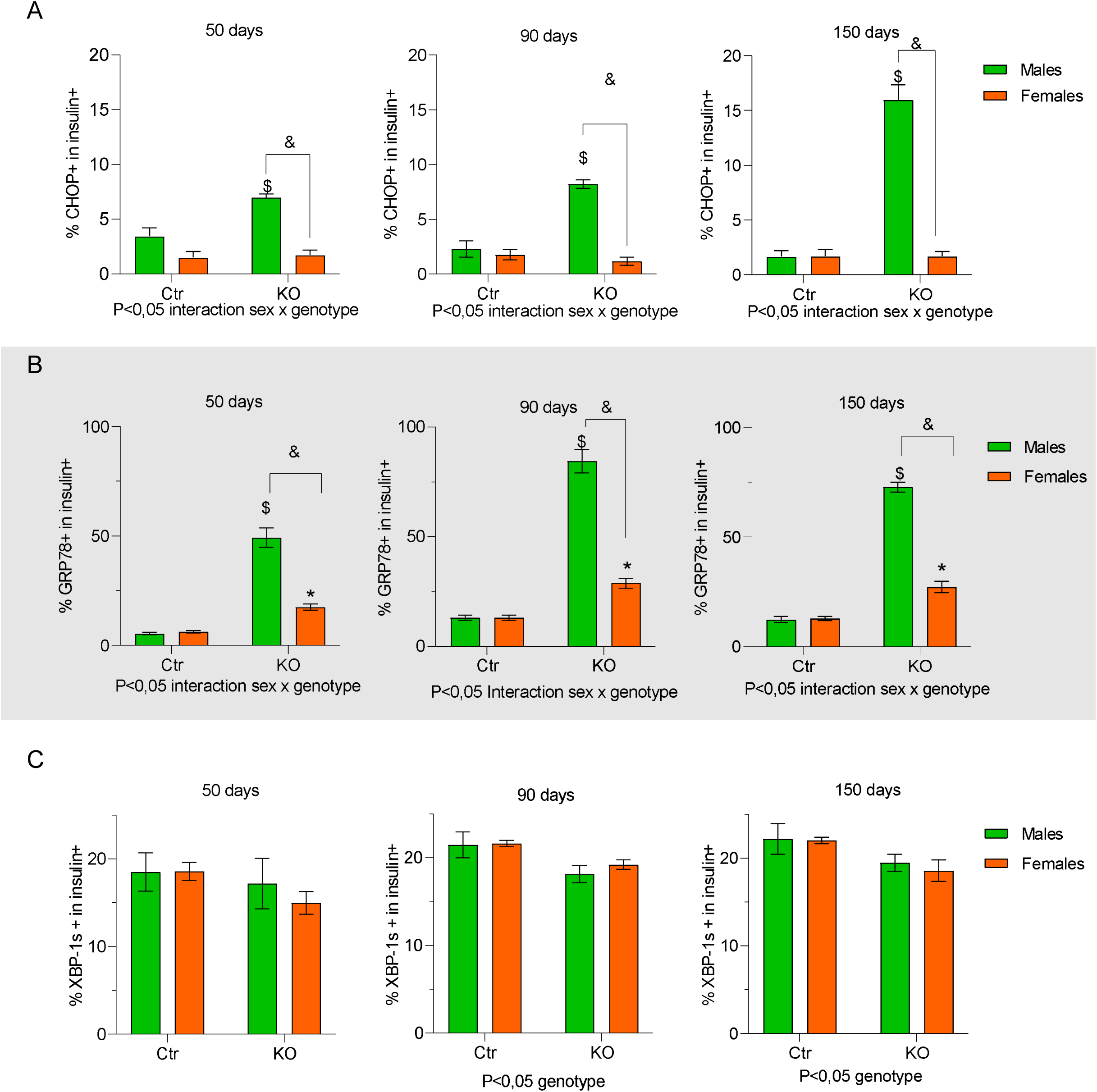
Quantification of the endoplasmic reticulum stress markers colocalized with insulin. Quantification of all 12 groups:4 genotypes FCtr (Female Control); MCtr (Male Control); FKO (Female KO) and MKO (Male KO) at 3 different ages, co-localization, expressed in positive percentual of either CHOP (A), GRP78 (B) or XBP1-s (C) positive cells in insulin+ cells. N=5 group, data is shown as mean± SEM, p<0,05: *FKO vs FCtr, $ MKO vs MCtr and & FKO vs MKO, source of variation is reported under the graph.

HNF4α KO also induces an increase of GRP78 expression on beta cells, observed by increased co-localization of GRP78 (red) and insulin (green) independent of gender (Figure 7B). However, at 50 days, the percentage of this co-localization was already higher in MKO than in FKO (Figure 8B). The increased expression of GRP78 in beta cells was progressive with age, being higher at 90 and 150 days in animals, and more significant for MKO than for FKO (Figure 7B and 8B). Thus, gender and age influenced HNF4α KO-induced GRP78 expression in beta cells.

Lastly, XBP-1s (red) co-localization with insulin (green) was similar in all groups evaluated (Figure 7 and 8C) at 50 days. However, in KO groups, there was a reduction on this co-localization, compared to Ctr after 90 days, independent of the gender (Figure 8C).

### Insulin-positive cells differentiated phenotype

Glut-2 (red) is co-localized with insulin (green) in all groups and ages analyzed (Figure 9A). However, after 90 days of age, FKO showed lower expression of Glut-2. This reduction of Glut-2 expression in insulin positive cells after 90 days of age was confirmed by quantification, and statistical analysis showed the influence of gender at 150 days of age (Figure 10A)

**Figure 9:**
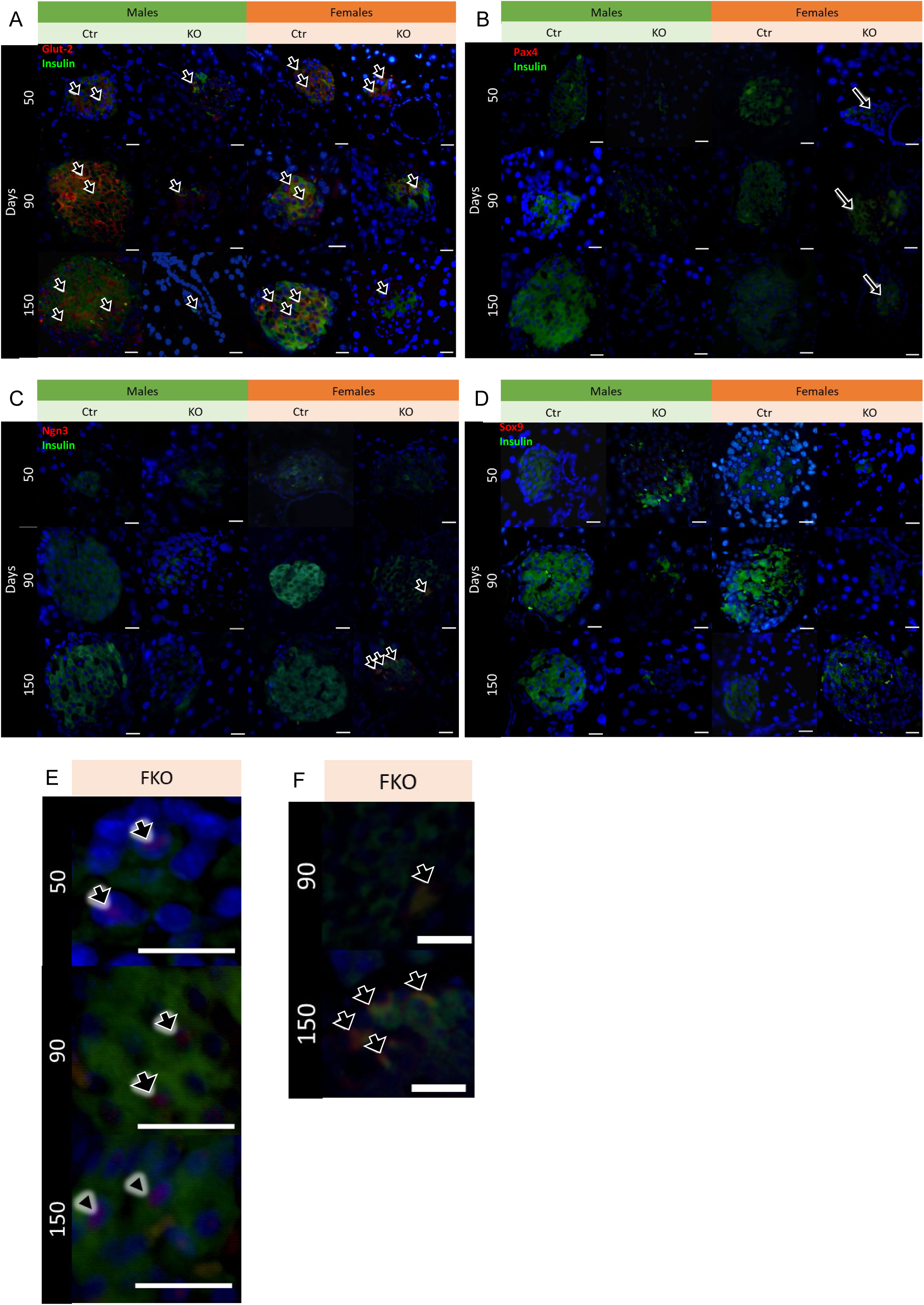
Beta cell phenotype markers colocalized with insulin. Representative images of all 12 groups:4 genotypes FCtr (Female Control); MCtr (Male Control); FKO (Female KO) and MKO (Male KO) at 3 different ages, immunofluorescence showing the co-localization of insulin (green) with either Glut-2 (red) (A), Pax-4 (red) (B), with highlights of the nuclear co-localization (purple) on the bottom right panel (E). NGN3 (red) (C), with highlighted co-localization on the bottom right panel (F) or SOX9 (red) (D) with insulin (green). Nuclei were stained with DAPI (blue). Scale bar: 50 μm.

**Figure 10:**
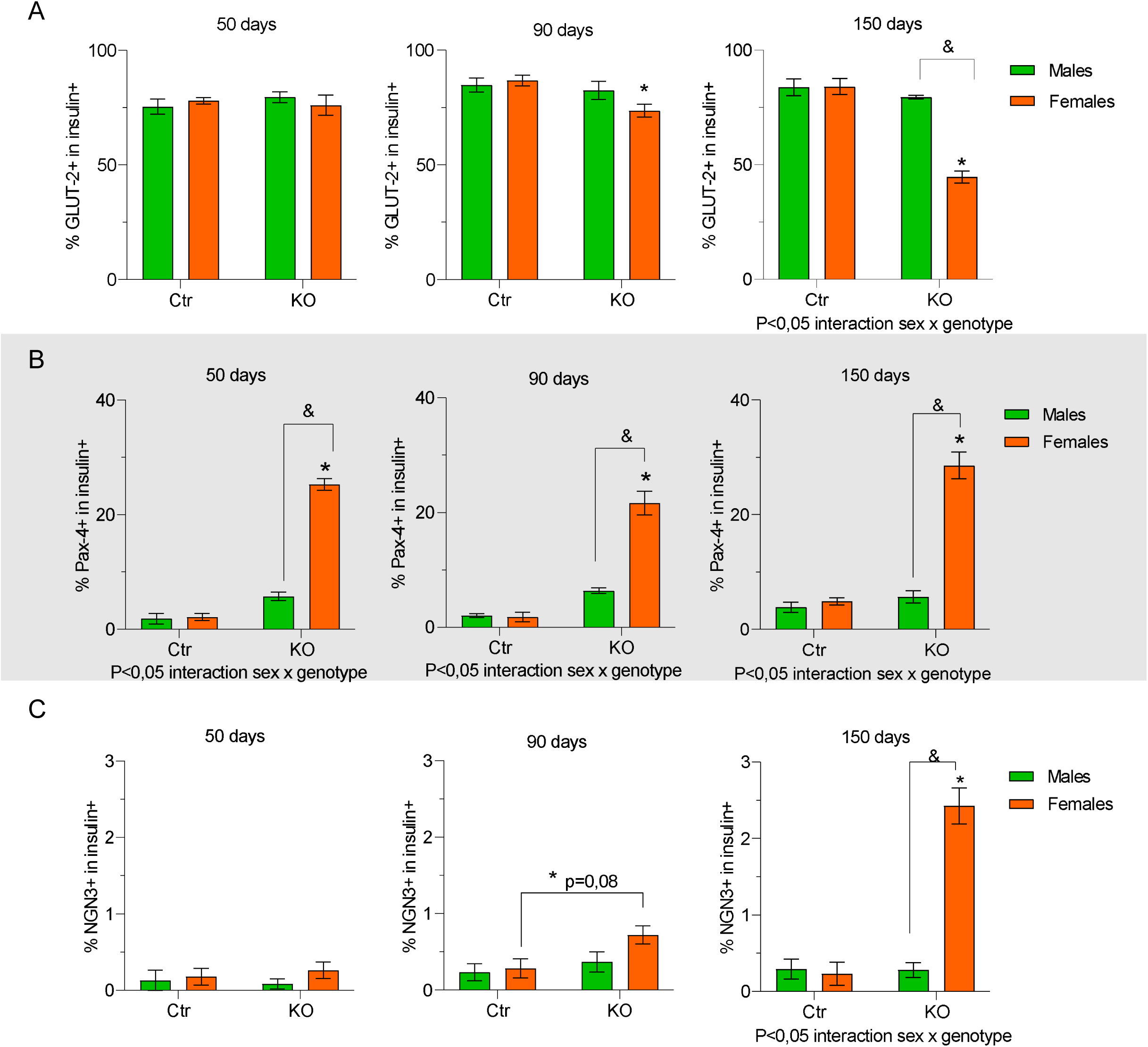
Quantification of the dedifferantiation markers colocalized with insulin. Quantification of all 12 groups:4 genotypes FCtr (Female Control); MCtr (Male Control); FKO (Female KO) and MKO (Male KO) at 3 different ages, co-localization, expressed in positive percentual of either Glut-2 (A), Pax-4 (B) or NGN3 (C) in insulin+ cells. N=5 group, data is shown as mean± SEM, p<0,05: *FKO vs FCtr, $ MKO vs MCtr and & FKO vs MKO, source of variation is reported under the graph., source of variation is reported under the graph.

Expressive Pax-4 (red) nuclear co-localization (purple) in insulin (green) positive cells is observed only in FKO animals, at all ages analyzed (Figure 9B, highlighted in 9E). Thus, this increased nuclear presence of Pax-4 in beta cells induced by HNF4α absence is influenced by gender (Figure 10B)

Similarly, increased NGN-3 (red) presence, which has a cytoplasmatic localization, in insulin (green) positive cells was observed only in FKO animals, at 90 and 150 days of age (Figure 9C, highlighted in 9F). However, this increase was only significantly higher in FKO compared to FCtr at 150 days of age. At that point, there was an influence of the gender, with MKO showing no alteration for NGN-3 expression (Figure 9C and 10C). We observed no expression of SOX-9 (red) in islet cells, immunostained with insulin (green), was similar for both genders and genotypes (Figure 9D).

## Discussion

The absence of HNF4α leads to impaired insulin secretion and loss of functional beta cell mass (Stoffel and Duncan 1997; Ellard 2000, 4; Gupta et al. 2005; 2007; Barth et al. 2022). MODY1 is a monogenic form of DM due to mutations in the transcription factor HNF4α with onset after puberty (Fajans 1989; Byrne et al. 1995, 1). Here, we have demonstrated for the first time that beta cell outcome after HNF4α absence is influenced not only by age but also by gender. Thus, although male and female mice with beta cell specific HNF4α knockout (MKO and FKO, respectively) become hyperglycemic, only the MKO have a worsening glucose intolerance dependent on age. We propose that the difference between genders in glucose handling loss can be attributed to different outcomes of UPR activation in the beta cells of our KO mice. Our data suggests that the MKO have a chronic activation of the UPR leading to apoptosis. On the other hand for FKO it does not progress to apoptosis, and instead induces loss of beta cell identity, an outcome of UPR activation previously described (Chen et al. 2022). In both cases, beta cell loss is exacerbated by age.

The rapid impairment of glucose tolerance due to HNF4α absence, observed here, is in line with its essential role in beta cell insulin production and secretion (Stoffel and Duncan 1997; Gupta et al. 2005). Interestingly, elevated fasting blood glucose a marker of chronic hyperglycemia progression (Wysham and Shubrook 2020), is only observed in MKO, suggesting that the beta cell loss observed in the males has a more significant functional impact.

Although previous work evaluating the role of HNF4α in beta cell function observed loss of glucose tolerance without changes to insulin positive cell percentage (Gupta et al. 2005), this study did not evaluate different ages and gender, which may explain the differences with our results. Further analysis of our KO mice islets revealed significant morphometric changes, with more expressive changes in circularity observed in MKO, presenting correlation with the more significant glucose tolerance dysfunction in this group, as it is observed that islet morphological changes may reflect a beta cell function change (Jacques-Silva et al. 2010). Indeed, the organization of cell types in the islet and its cytoarchitecture plays an essential role in beta cell function (Brereton et al. 2015a; Almaça, Caicedo, and Landsman 2020).

Alterations on islet morphology suggest that overall islet interactions may be disturbed, since the different islet cell types influence each other (Watts et al. 2016; Briant et al. 2018; Brereton et al. 2015). Thus, we evaluated alpha cells distribution in the islet, as it is known that beta cells can influence alpha cells proliferation and function (Mizukami et al. 2014; Watts et al. 2016). Indeed, we observed that impaired beta cell mass and insulin secretion resulted in increased percentage of glucagon positive cells per islet, which could further contribute to overall hyperglycemia.

To elucidate the mechanism through which HNF4α absence affects beta cell function, we evaluated expression of UPR components. We have shown that co-localization of GRP78 labelling and insulin is increased in the KO mice, with a notably higher and age dependent-increasing co-localization in MKO, correlating with glucose intolerance. Increased GRP78 expression is observed during ER stress in beta cell (Rutkowski and Kaufman 2004; Meyerovich et al. 2016; Cardozo et al. 2005). This suggested that absence of HNF4α leads to beta cell loss through altered UPR signaling. It has been shown that UPR signaling in beta cells has three outcomes: resolution of ER stress, apoptosis and loss of identity (Decio L. Eizirik and Cnop 2010; Chen et al. 2022). As such, we investigated these outcomes in our KO mice. Interestingly, only the MKO has increased CHOP co-localization with insulin-positive cells, indicating an activation of the apoptotic pathway in beta cells (Matsuda et al. 2010; Hu et al. 2019). Thus, it may be possible that progressive beta cell loss in MKO occur as a result of impaired beta cell ER function, due HNF4α absence that together with the progressive consequent hyperglycemia further aggravate the ER stress and beta cell loss via induction of ER apoptotic pathways (Décio L. Eizirik, Cardozo, and Cnop 2008; Decio L. Eizirik and Cnop 2010; Oslowski and Urano 2010).

Analysis of UPR pathway components demonstrates a gender-based difference in UPR activation outcome. Although there was reduced beta cell sXBP1, a marker of adaptative UPR, in both sexes, only in males, this was accompanied by increase in apoptotic marker and great induction of GRP78. This suggests that in females the UPR adaptive response seems to be impaired,, which may lead to loss of proper beta cell function (Herbert and Laybutt 2016; Chan et al. 2013) and loss of beta cell identity (Zhang et al. 2024; Chen et al. 2022). Indeed, proper sXBP1 expression is essential for maintaining beta cell identity (Lee et al. 2022) and other components of adaptative UPR (Back et al. 2009; Moin and Butler 2019; Chen et al. 2022).

One key marker of beta cell differentiation is Glut-2 expression (Neelankal John, Morahan, and Jiang 2017; Thorens 2015), which is reduced in FKO. In addition FKO beta cell shows an increase in PAX-4 expression at all ages analyzed, acute expression of this transcription factor is associated with a greater ability to reestablish ER homeostasis (Mellado-Gil et al. 2016), which may explain the lower activation of GRP78 expression in FKO beta cells. However, continuous expression of PAX-4 can lead to loss of insulin secretion, as it determines a less mature beta cells phenotype (He et al. 2011; Lorenzo et al. 2017). This is further highlighted by the presence of NGN3 positive cells in FKO, first observed at 90 days and more clearly at 150 days of age, as the reappearance of NGN3 positive cells in post-embryonic islets’ implies loss of fully mature beta cell state (Watada 2004; Talchai et al. 2012). The absence of SOX9 expression may indicate presence of cell with a phenotype between mature beta cells and endocrine progenitor (Kawaguchi 2013; Seymour 2014). Interestingly it has been described that beta cells that have this partially loss of identity can produce and secrete low volumes of insulin independently of changes in plasma glucose (Neelankal John, Morahan, and Jiang 2017; Baeyens et al. 2006). Therefore, a partial loss of identity in the beta cells of FKO mice could explain their better glucose tolerance than MKO, which needs further studies. These differences between genders are not related to the use of tamoxifen to induce HNF4α KO since we first confirmed that KO induction efficiency was similar to that previously described in the literature (Gupta et al. 2005) for both genders.

In conclusion, absence of HNF4α leads to impaired glucose tolerance and loss of insulin-producing beta cells, influenced by gender and age. Beta cell loss in KO groups may occur due to ER stress pathway alteration. Furthermore, it is possible to hypothesize that gender influences whether this dysfunction will be induced by the activation of pro-apoptotic pathways, as observed in male, or by beta cell dedifferentiation, as observed for female.

## Acknowledgments

The authors are grateful to Adriane Pereira Fernandes Araujo for her excellent technical assistance. We thank Drs. Antonio Boschero and Everardo Carneiro for donating the mice. We also thank the personnel from the Laboratory of Endocrine Pancreas and Metabolism (UNICAMP) and the technical personnel from the Department of Cell and Development Biology of ICB-USP.

---

**Supplementary Figure 1: HNF4α immunohistochemistry**. Representative images of all 12 groups (4 genotypes at 3 different ages), arrows point to some of the positive nuclei (A) grayscale of DAB filter (B), quantification of average positive nuclei per islet (C). N=5 group, data is shown as mean ± SEM. FCtr (Female Control); MCtr (Male Control); FKO (Female KO) and MKO (Male KO). p<0,05: *FKO vs FCtr, $ MKO vs MCtr and & FKO vs MKO, source of variation is reported under the graph Scale bar: 50 μm.

