## Supplementary figures and images for "The effect of HNF4α knockout in beta cells is age and gender dependent"

### Supplementary Figure 1

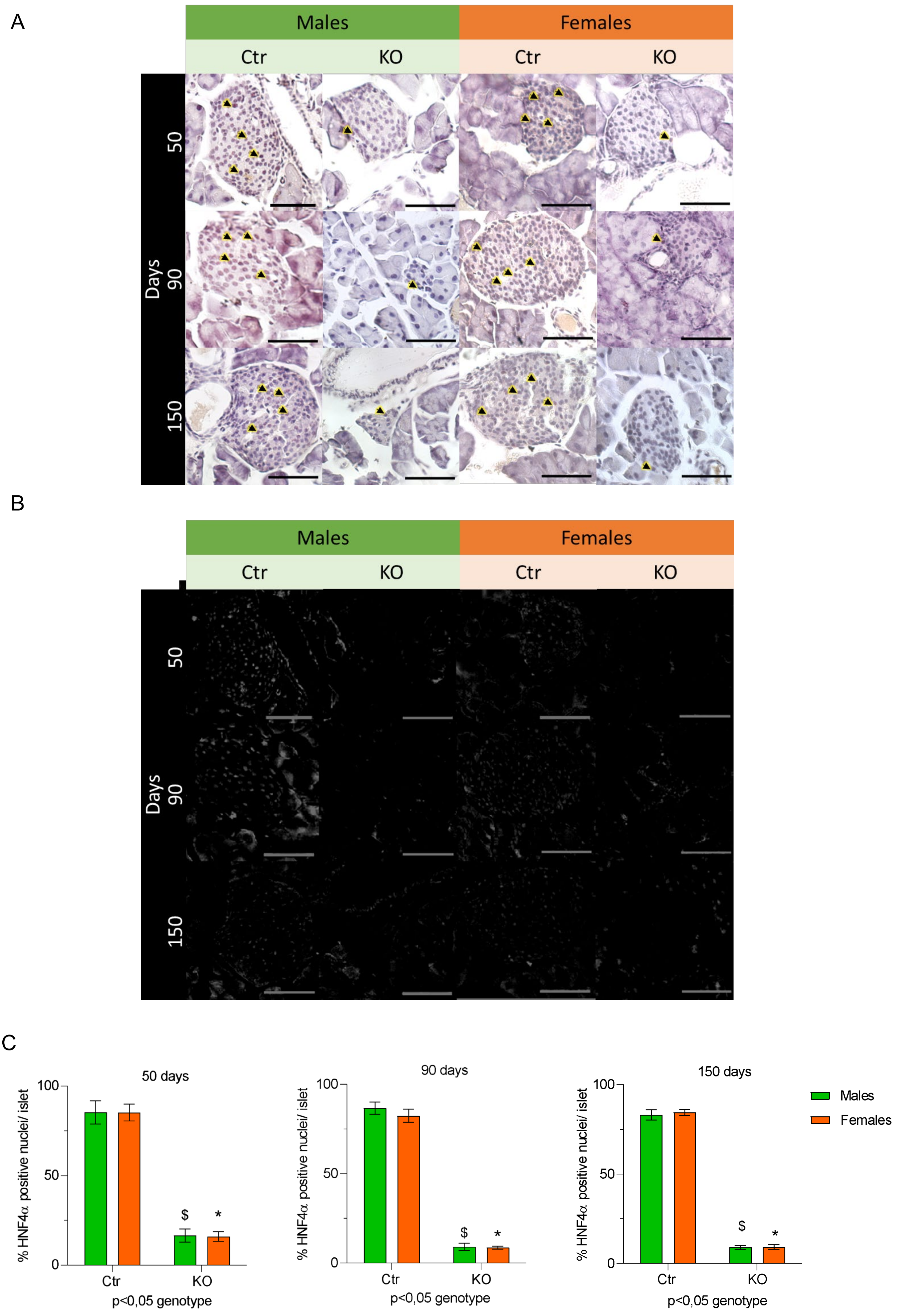
